# Evaluation of *in silico* algorithms for use with ACMG/AMP clinical variant interpretation guidelines

**DOI:** 10.1101/146100

**Authors:** Rajarshi Ghosh, Ninad Oak, Sharon E. Plon

## Abstract

**Background:** The American College of Medical Genetics and American College of Pathologists (ACMG/AMP) variant classification guidelines for clinical reporting are widely used in diagnostic laboratories for variant interpretation. The ACMG/AMP guidelines recommend complete concordance of predictions among all *in silico* algorithms used without specifying the number or types of algorithms. The subjective nature of this recommendation contributes to discordance of variant classification among clinical laboratories and prevents definitive classification of variants.

**Results:** Using 14,819 benign or pathogenic missense variants from the ClinVar database, we compared performance of 25 algorithms across datasets differing in distinct biological and technical variables. There was wide variability in concordance among different combinations of algorithms with particularly low concordance for benign variants. We also identify a previously unreported source of error in variant interpretation where *in silico* predictions are opposite to the evidence provided by other sources. We identified recently developed algorithms with high predictive power and robust to variables like disease mechanism, gene constraint and mode of inheritance, although poorer performing algorithms are more frequently used based on review of the clinical genetics literature (2011-2017).

**Conclusions:** Our analyses identify algorithms with high performance characteristics independent of underlying disease mechanisms. We describe combinations of algorithms with increased concordance that should improve *in silico* algorithm usage during assessment of clinically relevant variants using the ACMG/AMP guidelines.

## Background

Many *in silico* methods have been developed to predict whether amino acid substitutions result in disease. Use of this type of evidence has become a routine part of assessment of novel variants identified through gene-focused projects or as a part of whole exome or genome annotation pipelines. In a clinical setting, predictions from *in silico* algorithms are included as one of the eight evidence criteria recommended for variant interpretation by the American College of Medical Genetics and Genomics (ACMG) and Association of Molecular Pathologists (AMP)[1]. The ACMG/AMP guideline for use of *in silico* algorithms specifically states: “If all of the *in silico* programs tested agree on the prediction, then this evidence can be counted as supporting. If *in silico* predictions disagree, however, then this evidence should not be used in classifying a variant.” For a given missense variant, predictions by numerous algorithms are publicly available e.g. via dbNSFP[2] or Variant Effect Predictor[3] from which a few algorithms are typically chosen for variant interpretation and are often used without additional calibration. Different testing laboratories use distinct combinations of *in silico* algorithms for variant interpretation and this can lead to discordant interpretations. For example, in a recent assessment of the ACMG/AMP guidelines by the Clinical Sequence Exploratory Research consortium (CSER), the frequency of use of *in silico* algorithm evidence for pathogenic and benign variant assertion were 39% and 18% respectively[4]. The CSER study noted that use of *in silico* algorithms were one major source of discordance among different clinical laboratories and that the ACMG/AMP guideline for *in silico* algorithm usage may be aided by further recommendations[4].

Missense variants constitute a major set of variants of uncertain significance (VUS) in ClinVar[5]. An improved recommendation for use of *in silico* algorithms is important for reducing the VUS burden in clinical medicine and increasing concordance of variant interpretation. Currently, there is little consensus among clinical labs on how many and which algorithms to use for missense variant interpretation. For example, a recent exome sequencing study classified variants in 180 medically relevant genes for hereditary cancer according to ACMG/AMP guidelines. The authors found that the VUS rate was higher when requiring full concordance versus majority agreement among the 13 *in silico* algorithms used[6]. Other examples from literature demonstrate requiring full concordance among three[7] to seven[8] different algorithms for variant interpretation. However, to our knowledge, no analysis has been conducted to assess the applicability of the current ACMG/AMP guideline for *in silico* algorithm usage. Here, using predictions from 25 *in silico* algorithms for 14819 clinically relevant missense variants in the ClinVar database, we highlight several limitations of implementing the ACMG/AMP guideline for *in silico* algorithm usage. We find highly variable degree of concordance among different combinations of algorithms with particularly low concordance of the predictions of variants reported in ClinVar as benign. Using the ClinVar dataset, we identify algorithms with higher predictive power whose performances are robust to variables such as disease mechanism, level of constraint and mode of inheritance.

## Results

### Concordance among *in silico* algorithms

To identify the extent of concordance among *in silico* algorithms for known pathogenic and benign variants, we obtained 14,819 missense variants from ClinVar for which the rationale for pathogenic or benign assertion has been provided by at least one submitter (one star status in ClinVar), primarily clinical laboratories, and annotated these variants with scores and predictions from 25 algorithms using dbNSFP (v3.2)[9] or the respective authors’ websites. We generated a matrix of binary predictions (pathogenic or benign) for these variants with scores from the 18 algorithms, for which thresholds of pathogenicity were publicly available (Fig. 1A, Additional Table 1, see Methods). We found that when using this large number of algorithms that only 5.2% of the benign and 39.2% of pathogenic variants had concordant assertions across all them (Fig. 1A, Table 1). We obtained similar results when we restricted our analysis to benign and pathogenic variants in ClinVar that had identical assertions from at least two independent laboratories (two stars - Fig. 1B, Table 1) suggesting that errors in ClinVar assertions by a single submitter contributes little to the low level of concordance among algorithms. We then computed the pairwise differences among all the algorithms separately for 7346 benign and 7473 pathogenic variants in our dataset (see Methods). We found that on average, two algorithms tend to differ from each other significantly more in the interpretation of benign as opposed to pathogenic variants (p<0.0001, Welch Two Sample t-test) (Fig. 2A). Our data suggests that while interpreting large number of variants, full concordance, as suggested by the ACMG/AMP guidelines, is less likely to be achieved even when using only two algorithms, particularly for benign variants, consistent with earlier observation of poor correlations among predictors by Thusberg *et al* [10].

**Figure 1:**
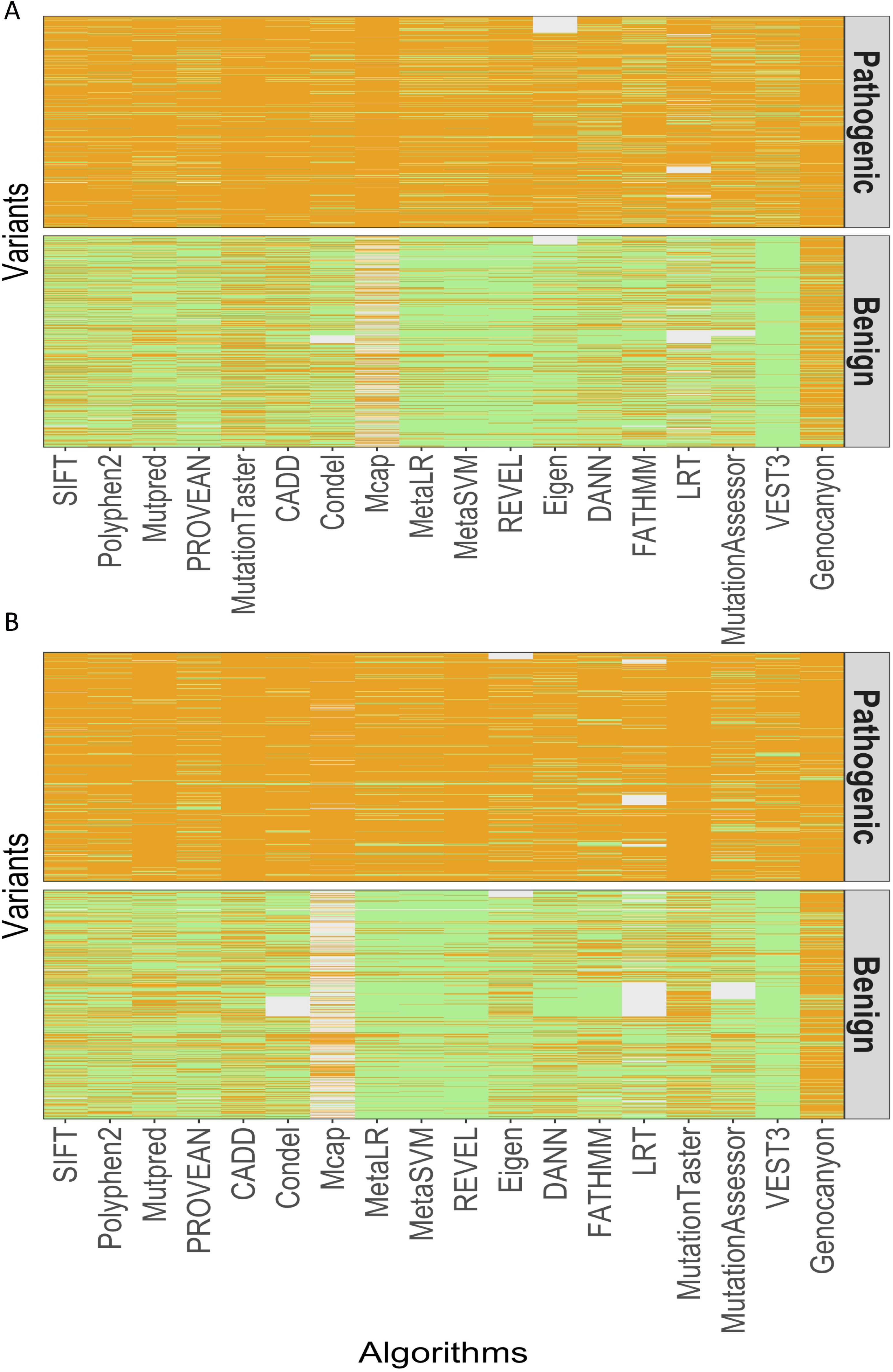
Concordance among predictions of 18 algorithms for variants in ClinVar. Binary predictions made by 18 algorithms for each pathogenic or benign variants in ClinVar are shown in upper and lower panel respectively. Each variant is along a row and an orange and a green tile depicts a pathogenic or benign call by the corresponding algorithm. 14819 variants with ClinVar review status one star or above (A) and 2966 variants with ClinVar review status two star or above (B) are shown.

**Table 1:** Concordance rate of different combination of algorithms

| Variant Assertion | Variants(n) | Concordance(%) | False concordance n(%) | Variant Source | Algorithms |
| --- | --- | --- | --- | --- | --- |
| Benign | 7346 | 5.2 | 57 (0.8) | ClinVar * | All 18 |
| Pathogenic | 7473 | 39.2 | 2(0.03) | ClinVar * | All 18 |
| Benign | 1914 | 4.5 | 12(0.6) | ClinVar ** | All 18 |
| Pathogenic | 1052 | 46.8 | 0(0) | ClinVar ** | All 18 |
| Benign | 7346 | 33.5 | 815(11.1) | ClinVar * | Polyphen;SIFT;CADD;PROVEAN;MutationTaster |
| Pathogenic | 7473 | 79.0 | 68(0.9) | ClinVar * | Polyphen;SIFT;CADD;PROVEAN;MutationTaster |
| Benign | 7346 | 46.2 | 1340(18.2) | ClinVar ** | Polyphen;SIFT;CADD |
| Pathogenic | 7473 | 84.9 | 156(2.1) | ClinVar ** | Polyphen;SIFT;CADD |

To assess the level of concordance among the most commonly used algorithms we reviewed algorithm use in the medical genetics literature between January 2011 and January 2017 (see Methods). We found that Polyphen[11] and SIFT[12] are cited most frequently followed by MutationTaster[13], CADD[14], PROVEAN[15], Mutpred[16] and Condel[17]. We did not detect any consistent pattern of combinations among these algorithms. In general, there seemed to be a bias in usage of some of the 25 algorithms while others, especially the more recently developed algorithms, are used less frequently. Predictions from five commonly used algorithms (Polyphen, SIFT, CADD, PROVEAN and MutationTaster) resulted in higher concordance relative to all 18 algorithms, but with 79% for pathogenic variants and only 33% for benign variants (Table 1).

In addition to lack of full concordance in prediction, we identified 815 of 7346 (11.1%) benign variants in ClinVar for which all five commonly used algorithms predicted the variant to be pathogenic and conversely 68 of 7473 (0.9%) pathogenic variants in ClinVar were predicted benign by all algorithms, referred to here as false concordances (Table 1). In fact, 22.5% (1653/7346) of benign variants were assessed as pathogenic by the 50% or more of the algorithms including 87 variants where the benign classification of the variant had been reviewed by a ClinVar recognized expert panel (3-star review status) suggesting that these are benign and not misclassified variants. In comparison, 5.2% (389/7473) of ClinVar pathogenic variants were deemed to be benign by the majority of algorithms (Additional Table 2). Evaluating only three commonly used algorithms (Polyphen, SIFT and CADD) resulted in higher concordance for pathogenic (84%) and benign (46%) variants, however, coupled with an increase in false concordances (Table 1).

**Table 2:**
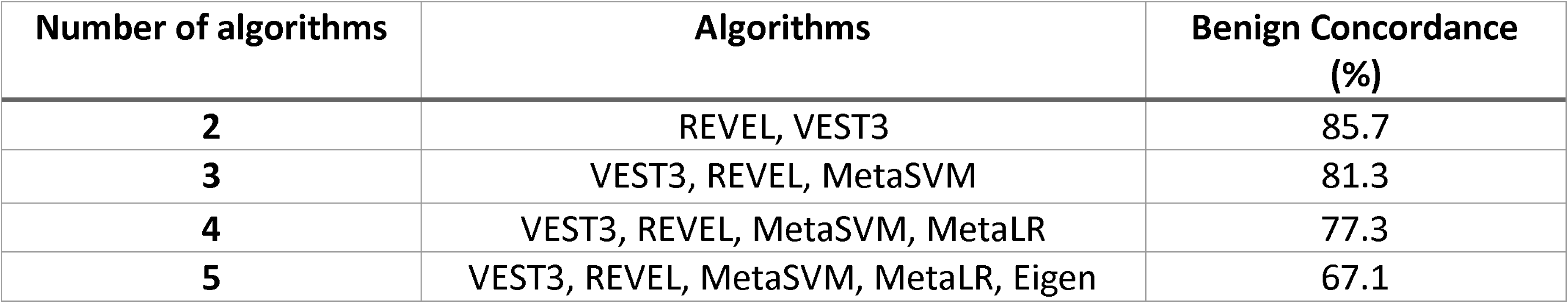
Percentage of variants with concordant benign assertions.

| Number of algorithms | Algorithms | Benign Concordance (%) |
| --- | --- | --- |
| **2** | REVEL, VEST3 | 85.7 |
| **3** | VEST3, REVEL, MetaSVM | 81.3 |
| **4** | VEST3, REVEL, MetaSVM, MetaLR | 77.3 |
| **5** | VEST3, REVEL, MetaSVM, MetaLR, Eigen | 67.1 |

Not surprisingly, we failed to identify any combinations of algorithms that resulted in false concordance of zero and true concordance of 100% among the 18 algorithms whose default predictions are publicly available. We generated all possible combinations of three (n=816), four (n= 3060) or five (n=8568) algorithms and obtained their true and false concordance rates across the 14,819 variants. As before, there was a lower false concordance rate and a higher true concordance rate for pathogenic variants relative to the benign variants (Fig. 2B). Overall, the concordance among combinations for benign variants ranged from 85% to 67% for a pair to five algorithms, respectively (Table 2). We noted that the best performing combinations of algorithms were different for benign and pathogenic variants (Additional Table 3). For example, for benign variants the best performing combinations of three algorithms consisted of VEST3[18], REVEL[19] and MetaSVM [20] with a true concordance rate of 81.3% and a false concordance rate of 2.8 %, whereas for pathogenic variants the same combination resulted in a 70% true concordance and a 5.4 % false concordance. For pathogenic variants, the best performing trio combination consisted of MutationTaster, Mcap[21] and CADD (Additional Table 3). We obtained similar results for combinations of four or five algorithms (Additional Table 3). In general, many different combinations performed better for pathogenic than benign variants (Fig. 2B).

**Figure 2:**
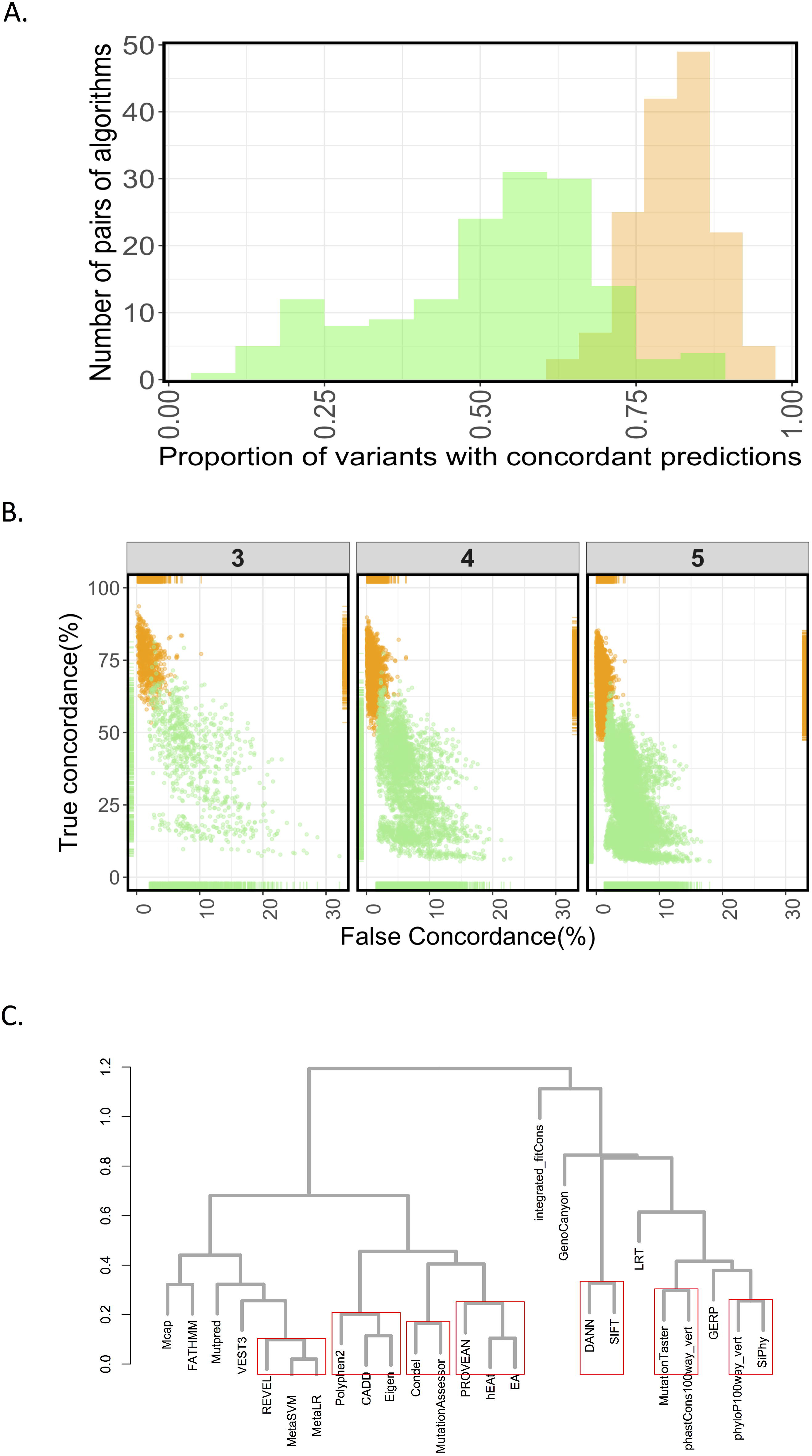
Concordance among algorithms. A) Distribution of proportion of variants that had concordant calls by any given pair of algorithms (among 18 algorithms) for benign (green) and pathogenic (orange) variants in ClinVar. B) Scatterplots of true concordance (variant assertion matches ClinVar assertion) vs False Concordance (Variant assertion does not match ClinVar assertion) for combinations of 3, 4 or 5 algorithms at a time. An orange and a green point depicts the true and false concordance of a combination for benign and pathogenic variants, respectively, in ClinVar. C) Hierarchical clustering of 25 algorithms with scores for 14819 variants in ClinVar. Red rectangles indicate robust clusters with an AU p-value of >0.99 (see methods).

Taken together, our results suggest that a given combination of algorithms (using the publicly available threshold scores) will perform quite differently across benign and pathogenic variants with a significant chance of erroneous assertion due to false concordance among algorithms. These false concordances could potentially bias the variant interpretation towards a VUS classification if all the other available variant data suggests the opposite assertion.

### Further analysis of algorithm prediction and concordance

For some algorithms such as Eigen[22], hEAt [23], GERP[24] etc. cut-offs defining pathogenic or benign assertion are either not recommended or inferred arbitrarily. We therefore used the actual output scores provided by all 25 algorithms as a continuous variable to identify algorithms whose predictions are likely to be concordant independent of the algorithms internal cut-offs. A hierarchical clustering of the normalized output scores of 14,819 missense variants for each of the algorithms revealed seven clusters (Fig. 2C). All the largely evolutionary conservation algorithms such as phyloP[25], phastCons[26], GERP[27] and Siphy[28] belong to different clusters from the metapredictors REVEL, MetaSVM and MetaLR (Fig. 2C).

### Comparison of performance of *in silico* algorithms

To identify well-performing algorithms with prediction abilities that are robust to the nature of a variant, gene constraint and underlying disease mechanism, we quantified performance of the *in silico* algorithms on multiple test datasets by determining the area under the receiver operator characteristic curve (AUC) (see Methods).

We analyzed two overlapping datasets differing in the confidence of variant assertions. These were 14,819 ClinVar variants that are assigned at least one star review status and 2966 ClinVar variants with concordant scores from at least two laboratories (two stars - see Methods). For both datasets, we observed wide variation in performance of the algorithms with AUCs ranging from 0.5 to 0.96 (Fig. 3A). We identified several algorithms with AUC ≥0.9 in these datasets that did not differ significantly in their performance between ≥1 or ≥2 star datasets (Fig. 3A).

**Figure 3:**
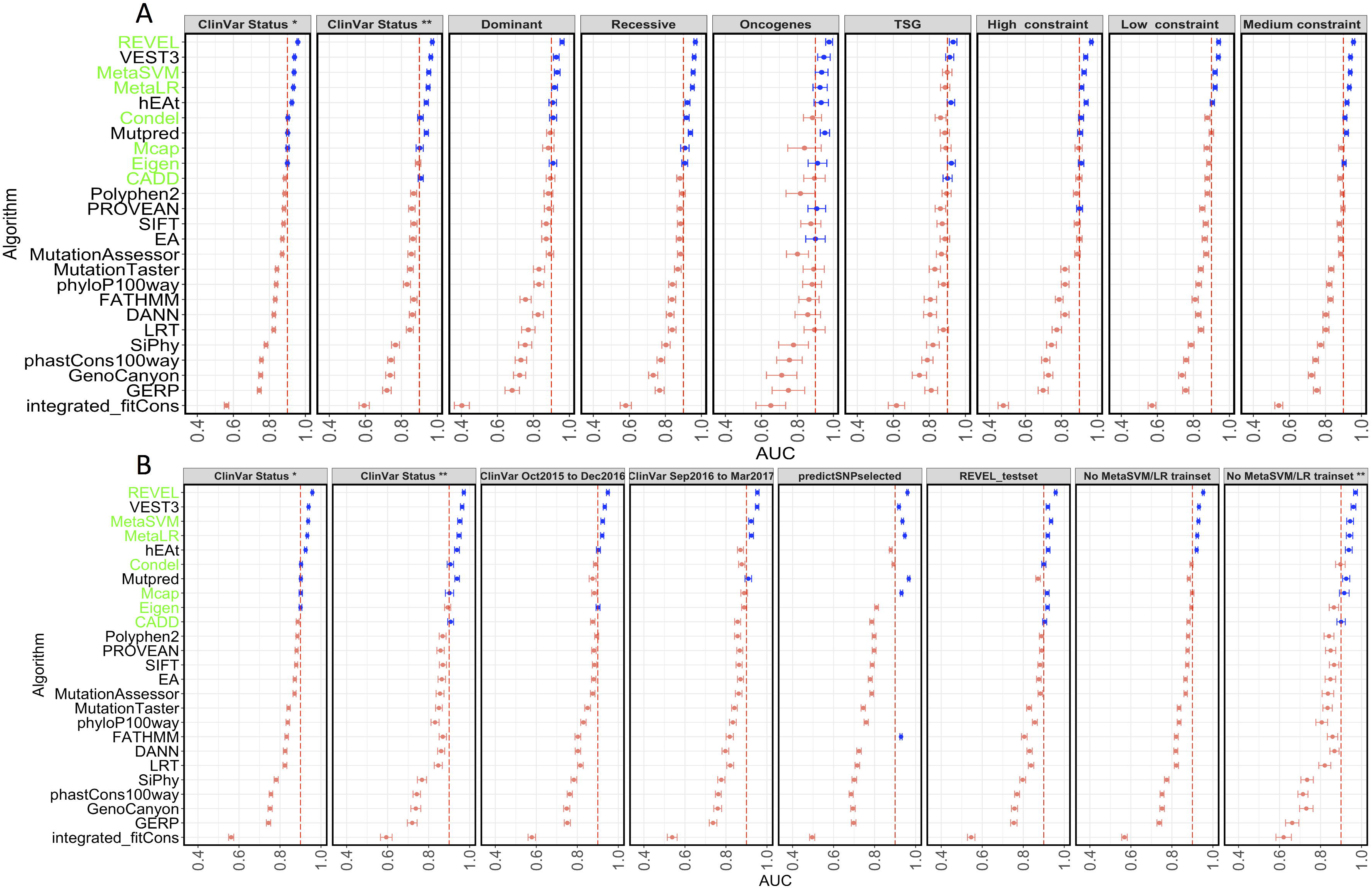
Performance analysis of algorithms. The AUC of a ROC are plotted for 25 algorithms. Vertical dotted line indicates a AUC of 0.9 and 99% confidence intervals for each AUC are shown. Blue dots indicate AUC>0.89. A) AUCs of the algorithms across five different datasets shown in the panels and described in text. B) AUCs of the algorithms across five different datasets (represented in panels) shown in the panels to address type I circularity as described in text.

We next sought to identify algorithms whose performance did not differ whether a given variant resulted in gain-of-function (GOF) or loss of function (LOF) of a gene product by analyzing datasets enriched in activating/GOF mutations in oncogenes and LOF mutations in TSG which are both pathogenic in cancer development[29] (see Methods). We also separately evaluated 1169 benign and 1427 pathogenic variants in genes linked to diseases with primarily recessive mode of inheritance as another proxy for a dataset enriched in LOF variants. We did not observe significant differences in performance of algorithms in the GOF and LOF datasets including across the high performing algorithms (Fig. 3A). Additionally, we analyzed variants in genes that are primarily linked to diseases with dominant mode of inheritance. The latter dataset is likely a mixture of LOF and GOF variants. Again, there was no major departure from the rank-order of the top five performing algorithms that we observed in the other datasets (Fig. 3A).

Finally we explored whether the performance of algorithms were affected by the level of constraint on a gene, as defined by the comparing the expected and observed missense variants in ExAC[30] (missense Z scores). We obtained variants in genes with high, intermediate or low level of constraint by a missense Z score threshold of >2.5, between 0 to 2.49 or less than zero respectively. We did not observe any major changes in the rank-order of the algorithms (Fig. 3A).

Taken together our analyses suggests that the performance of the majority of algorithms in current use are unlikely to be affected significantly as a function of the nature of a variant or the level of constraint on the gene. The high-performing algorithms are robust to these variables and are more likely to give rise to consistent interpretation across different variant datasets.

### Evaluation of potential circularity in algorithm analysis

There is significant concern that the result of analyses such as that described here may arise from circularity in data used. For example, REVEL, a meta-predictor whose features includes 13 out of the 25 algorithms that we analyzed[19], performed best in all nine datasets described above. It was possible that the variants we used to assess performance of REVEL and other algorithms described here were also included in its training sets, which for REVEL included some of the ClinVar and HGMD variants available until October 2015. This type of circularity inflates the performance measures of some algorithms and is referred to as type I circularity[31]. To examine the possibility of type 1 circularity inflating the performance of algorithms that were trained on HGMD and ClinVar variants, we compared performance of all the algorithms in six additional datasets: A) ClinVar Oct2015 to Dec2016: This dataset consists of ClinVar missense variants with ≥1-star review status that were released between October 2015 and December 2016. B) ClinVar Sept 2016 to March 2017: This even more recent set of missense ClinVar variants with ≥1 star is absent in ClinVar data releases prior to September 2016. The A and B datasets consists of newer variants that are likely to be absent in the training sets of algorithms that were developed earlier. In addition, the newer ClinVar variants are also more likely to have been classified using ACMG/AMP 2015 guidelines, which recommends only “supporting” weight for *in silico* evidence towards pathogenicity classification. Thus, it is likely that the clinical laboratory primarily relied on independent clinical and genetic data to come to the final variant assertion. C) predictSNPselected[31] is a benchmark dataset that does not contain the CADD training data. D) REVEL test set exclude the variants in HGMD and ClinVar that were used for training REVEL[19]. E) and F) Minus MetaSVM/LR trainset: We removed all the variants that were used in training the metapredictors MetaSVM and MetaLR[20] from our ClinVar variant set. These datasets consisted of variants designated ≥1-star (E) or ≥2-star(F) review status in ClinVar. The resulting predictions of these datasets which removed variants used in training different algorithms did not demonstrate a major change in the rank order of the top five algorithms that exhibited an AUC>0.9 with REVEL performing the best in these datasets (Fig. 3B, Additional Fig. 1).

We next tested if the top performing algorithms suffered from type 2 circularity[31], which has been described as a caveat introduced due to the reliance of an algorithm’s performance on the distribution of pathogenic or benign variants in a protein. Thus, in a variant dataset where there are proteins with only pathogenic variants or only benign variants (unbalanced dataset) some algorithms tend to perform better than in a dataset that have equal number of pathogenic and benign variants per gene (perfectly balanced), even if this is not what is biologically present. To this end we compared performance on an unbalanced dataset (Varibenchselected[31]) and a balanced dataset which includes equal number of pathogenic and benign variants per protein in ClinVar (see Methods). Consistent with earlier results[31] we found that FATHMM[32] is particularly sensitive to this type of circularity. In other words, there is drop in performance of FATHMM in analysis of a dataset that is perfectly balanced (Balanced dataset * in Additional Fig. 2). We also detected evidence for potential type 2 circularity for algorithms such as MetaSVM/LR and MCAP (Balanced vs varibench for MetaSVM, MetaLR and MCAP, bootstrap p value <0.0001, Additional Fig. 2). Thus, caution should be used in interpreting scores using algorithms such as FATHMM as the prediction efficacy is partly dependent on pathogenic and benign variant distributions in any given gene.

### Discussion

The ACMG/AMP guideline for use of *in silico* algorithms in a clinical setting suggests full concordance among multiple algorithms for this type of evidence to be used in missense variant classification without further clarification of the number or choice of algorithms. As we have shown, such usage leads to discrepancies arising mainly because of the lack of specification. Our review of the literature reveals that the metric for concordance is not consistent across different laboratories. While some studies have adhered to the ACMG/AMP guidelines for strict concordance, others have used a majority vote rule. It has been reported that use of the strict ACMG criteria gave rise to a higher rate of VUS[6] and increased discrepancies among laboratory classifications[4]. The lack of a standard guidance for incorporating *in silico* algorithms could potentially lead to increased VUS burden and inter-lab discrepancies.

In addition, we find that frequently used algorithms are older and vary in performance. Our analyses identified several high performance relatively newer algorithms which are infrequently used such as REVEL, VEST3 etc. Many of these algorithms are ensemble predictors incorporating many older algorithms as features. The performances of these algorithms are robust to technical artifacts, levels of constraint on genes, the underlying nature of variants and Mendelian inheritance pattern. Thus, laboratories may benefit from modifying pre-existing variant interpretation pipelines that currently use older algorithms.

The ACMG/AMP guideline encourages use of multiple algorithms. Conversely, we observed an increase in the discordant calls as more algorithms are used to infer variant pathogenicity thus hindering the use of *in silico* evidence. An alternative is to use metapredictors that in effect combine multiple individual predictors to generate a score. These metapredictors satisfy the concept underlying the multiple algorithm criteria; of note, combining them with their constituent predictors for variant interpretation may not be ideal.

In general, we show, using author recommended thresholds for variant assertion, a substantial increase in likelihood of discordance, particularly for benign variants. We found that for pathogenic variants, the concordance among algorithms were higher most likely due to the tendency of several algorithms to call a variant pathogenic leading to incorrect inferences. Consistent with this, we found several variants for which multiple algorithms made concordant assertions that were opposite to what is reported in ClinVar. Although this could be a result of misclassification in ClinVar, we found that a ClinVar designated expert panel has interpreted some of these variants. These false concordances are another source of error for variant interpretation. The problem of false concordance both increases the VUS rate and highlights why it may be inappropriate to increase the ACMG/AMP evidence strength for computational algorithms from “supporting” to “moderate” or “strong”. We independently identified combinations of algorithms that tend to be more concordant via a hierarchical clustering of the output scores of all algorithms. The clustering pattern suggested that it is probably best to make inferences separately for evolutionary conservation algorithms e.g. GERP and metapredictors. Combining them is likely to result in discordant calls. Another alternative may be to calibrate the thresholds with known variants in genes under consideration.

Our results are not designed to identify a single algorithm for use across all genes although data suggests that high performing algorithms perform well across many different gene and mutation mechanism type. In addition, gene specific algorithms or gene specific calibration of algorithms using well-characterized set of benign and pathogenic variants may perform better than the general approach described here. We note that several algorithms are very sensitive to the multiple sequence alignment[33]. The performance of SIFT and other algorithms within our analyses and others such as Align-GVGD[34] could potentially be improved if gene-specific curated alignments are provided.

### Conclusions

The analyses and the data presented in this article highlights problems associated with the strict use of ACMG /AMP guidelines for *in silico* algorithm usage. In particular our results identify poor concordance among algorithms, particularly for benign variants. We highlight the problem that the concordance rate vary substantially depending on the combination of algorithms, which contribute to inter clinical laboratory discrepancy in variant assertion. We also identify a previously unreported source of error in variant interpretation where *in silico* predictions are opposite to the evidence provided by other sources (false concordance). Finally, we identify and provide high performing algorithm combinations many of which are recently developed. Taken together, this analysis provides the necessary data and framework for optimization of the ACMG guidelines and offers methods to potentially reduce the burden of variants of uncertain significance in clinical variant interpretation.

## Methods

### Code and data availability

All the data necessary to produce the figures in the manuscript and the associated code are included as Additional data. The code is also available here: http://rpubs.com/thisisrg/supp_code_17

### Variant data and annotation

We downloaded the variant_summary.txt files from the ClinVar ftp site for variant used in the analysis. In this manuscript we used the files <u>http://ftp.ncbi.nlm.nih.gov/pub/clinvar//tab_delimited/archive/variant_summary_2016-09.txt.gz</u> and <u>http://ftp.ncbi.nlm.nih.gov/pub/clinvar//tab_delimited/archive/variant_summary_2016-12.txt.gz</u> along with their corresponding xml files. We removed all variants whose review status were “no assertion criteria provided”. We also excluded any variants of uncertain significance from our analysis. We next considered only the missense variants and filtered out all the other classes of variants such as frameshift, termination, silent, non-coding etc. Finally, we collapsed the Likely pathogenic and Pathogenic variants in one group and Likely Benign and Benign variants in another group. Thus, our final data of 14819 variants had two levels of clinical significance: Pathogenic and Benign (Additional data 1).

### Algorithms and scores

We annotated these variants with 25 algorithm scores using dbNSFP and authors’ publicly available websites (Additional table 1). To generate binary predictions, we used the threshold recommended by dbNSFP3.2 or by the algorithms’ authors. Certain algorithms such as MutationTaster, Mutation Assessor, and Polyphen have thresholds such that it generates more than two classes. We collapsed the “probably damaging” and “possibly damaging” classes variants of Polyphen into a single “damaging” class. For MutationTaster we collapsed the “A” (disease causing automatic) and “D” (disease causing) classes into a single “damaging” class, while the "N" ("polymorphism") or "P" ("polymorphism_automatic") were collapsed into a single “Tolerated” class. For MutationAssessor that generates four predictions, the high ("H") or medium ("M") categories were treated as “Damaging” whereas the low ("L") or neutral ("N") categories were treated as “Tolerant”. LRT predictions in dbNSFP gives three classes namely “Damaging”, “Neutral” and “Unknown”. We treated the “Unknown” labels as no data available or NA in our analysis. For certain algorithms such as SIFT, MutationTaster, PROVEAN and FATHMM multiple scores for a given variant corresponding to different transcripts are provided by dbNSFP. We used the most damaging score predicted by the corresponding algorithm for a given variant in our analyses.

### Literature search

To identify the frequency of usage of algorithm from the year 2011 to 2017 we conducted a literature search in PubMed (search date Jan 19, 2017) using PubmedReminer (<u>http://hgserver2.amc.nl/cgi-bin/miner/miner2.cgi</u>) using the following search string: "humans"[MeSH Terms] AND Medical Genetics[filter] AND ("SOMATIC"[ALL FIELDS] OR MISSENSE[ALL FIELDS] OR GERMLINE[ALL FIELDS] OR ("mutation"[MeSH Terms] OR "mutation"[All Fields]) OR VARIANT[All Fields] OR ("polymorphism, genetic"[MeSH Terms] OR ("polymorphism"[All Fields] AND "genetic"[All Fields]) OR "genetic polymorphism"[All Fields] OR "polymorphism"[All Fields]) AND (dbnsfp[all fields] OR POLYPHEN[ALL FIELDS] OR SIFT[ALL FIELDS] OR VEST3[All Fields] OR METASVM[ALL FIELDS] OR METALR[ALL FIELDS] OR CONDEL[ALL FIELDS] OR CADD[ALL FIELDS] OR MUTATIONASSESSOR[ALL FIELDS] OR PROVEAN[ALL FIELDS] OR FATHMM[ALL FIELDS] OR EIGEN[ALL FIELDS] OR MUTPRED[ALL FIELDS] OR "REVEL"[ALL FIELDS] OR DANN[All Fields] OR LRT[All Fields] OR MUTATIONTASTER[All Fields] OR GERP[All Fields] OR VEST3[All Fields] OR Genocanyon[All Fields] OR fitcons[All Fields] OR phastcons[All Fields] OR phylop[All Fields]) AND ("2011/01/01"[PDAT]: "2017/12/31"[PDAT]) NOT (18570327[UID] OR 19734154[UID] OR 20052762[UID] OR 20642364[UID] OR 23990819[UID] OR 22077404[UID] OR 21763417[UID] OR 21457909[UID] OR 21480434[UID] OR 21412949[UID] OR 23033316[UID] OR 22949387[UID] OR 22689647[UID] OR 22539353[UID] OR 27577208[uid] OR 27468419[uid] OR 27357839[uid] OR 27224906[uid] OR 27148939[uid] OR 27147307[uid] OR 27128317[uid] OR 23620363[UID] OR 23315928[UID] OR 27841654[uid] OR 27721395[uid] OR 27760515[uid] OR 27564391[uid] OR 27995669[uid] OR 24487276[UID] OR 24205039[UID] OR 25073475[UID] OR 25684150[UID] OR 26555599[uid] OR 27776117[UID] OR 26426897[uid] OR 26332131[uid] OR 27666373[UID] OR 26982818[uid] OR 26892727[uid] OR 26885647[uid] OR 26866982[uid] OR 26727659[uid] OR 26681807[uid] OR 26633127[uid] OR 26677587[uid] OR 26504140[uid] OR 26269570[uid] OR 26015273[uid] OR 24675868[uid] OR 24648498[uid] OR 24651380[uid] OR 24453961[uid] OR 24451234[uid] OR 24338390[uid] OR 24332798[uid] OR 25979475[uid] OR 25967940[uid] OR 25851949[uid] OR 25599402[uid] OR 25587040[uid] OR 25557438[uid] OR 25552646[uid] OR 25535243[uid] OR 25519157[uid] OR 25393880[uid] OR 23020801[uid] OR 22937107[uid] OR 22747632[uid] OR 22322200[uid] OR 22261837[uid] OR 22110703[uid] OR 22192860[uid] OR 21925936[uid] OR 21919745[uid] OR 21814563[uid] OR 25117149[uid] OR 24980617[uid] OR 24718290[uid] OR 24194902[uid] OR 23954162[uid] OR 23935863[uid] OR 23819846[uid] OR 23843252[uid] OR 23836555[uid] OR 23462317[uid] OR 23424143[uid] OR 23357174[uid] OR 21685056[uid] OR 21520341[uid] OR 20866645[uid] OR 20689580[uid] OR 20625116[uid] OR 20084173[uid] OR 19602639[uid] OR 19105187[uid] OR 18990770[uid] OR 18654622[uid] OR 18325082[uid] OR 18384978[uid] OR 18195713[uid] OR 18186470[uid] OR 18179889[uid] OR 18005451[uid] OR 17989069[uid] OR 17537827[uid] OR 27058395[uid] OR 26567478[uid] OR 26095143[uid] OR 22997091[uid] OR 22038522[uid] OR 20660939[uid] OR 20224765[uid] OR 19217021[uid] OR 18361419[uid] OR 18210157[uid] OR 17349045[uid] OR "REVIEW"[PUBLICATION TYPE] OR "REVIEW LITERATURE AS TOPIC"[MESH TERMS] OR REVEL[AU] OR DANN[AU] OR 26566084[uid] OR 26328548[uid] OR 26054510[uid] OR 24369116[uid] OR 23824587[uid] OR 22974711[uid] OR 20717976[uid] OR 20613780[uid] OR 18797516[uid] OR 23223146[uid] OR 26025364[uid] OR 26961892[uid] OR 26098940[uid] OR 25878120[uid] OR 25340732[uid] OR 24740809[uid] OR 24442417[uid] OR 24266904[uid] OR 24065196[uid] OR 24037343[uid] OR 23571404[uid] OR 23148107[uid] OR 21827660[uid] OR 21536091[uid] OR 21107268[uid] OR 19648217[uid] OR 19116934[uid] OR 18615156[uid] OR 18463975[uid] OR 18252211[uid] OR 18161052[uid] OR 24482837[uid] OR 23274505[uid] OR 22940547[uid] OR 22912676[uid] OR 21575667[uid] OR 19786005[uid] OR 19562469[uid] OR 19444471[uid] OR 19255159[uid] OR 19142206[uid] OR 19138047[uid] OR 18991109[uid] OR 18602337[uid] OR 18552399[uid] OR 18541031[uid] OR 18357615[uid] OR 18203168[uid] OR 17722232[uid] OR 17456336[uid] OR 17431481[uid] OR 17375033[uid] OR 17375033[uid] OR 17375033[uid] OR 17375033[uid] OR 28093075[uid])

Briefly we restricted our analysis to the medical genetics literature and excluded reviews and technical papers reporting discovery and comparative analysis of algorithms as defined by the above search term. We obtained 507 of articles that mentioned an algorithm in the Title or abstract. The number of articles per algorithm term was used as a proxy for the usage of algorithms used in our analysis.

### Concordance analysis

To determine concordance among algorithms we obtained the publicly available thresholds (Additional table 1) to define a dataset of pathogenic and benign prediction for each variant. We next generated all possible pairwise combinations of algorithms and determined the proportion of variants for which they agree with each other. Next we also generated all possible combinations of algorithms with 3, 4 or 5 members and determined the concordance with ClinVar assertions for each of these pairs. We also determined the fraction of variants for which algorithms in each combination was concordant but the assertion was opposite to that designated in ClinVar. We refer to these instances as false concordances. A list of such combinations and their true and false concordances are provided in the Additional data 2-6.

### Clustering

The scores for 25 algorithms for each of the 14,819 variants were used to cluster the algorithms using pvclust package in the R programming environment. We identified the most confident clusters by using 50000 bootstrap replicates of the data, Euclidean distance as a measure of similarity and ward’s D2 method of hierarchical clustering as implemented in the pvclust function[35]. We called clusters as stable if they had a 0.99 or above probability of having the same members in the bootstrap replications. The final rendering of the plot was done using the dendextend[36] package in R.

### Performance analysis

We compared the performance of each of the algorithms on all datasets separately by estimating the area under the curve (AUC) of a receiver operator characteristic (ROC) curve and its 99% confidence interval using the OptimalCutpoints library in R. We estimated significant differences between any two AUCs by using 10000 stratified bootstrap replicates of the datasets in question (where each replicate contained the same number of benign and controls than in the original sample), calculating AUC for each replicate for each and then testing for the statistical significance as implemented in the library pROC in R.

### Datasets

All the data are available as additional data files or are available from the respective authors. We provide brief descriptions of the datasets that we used below:

ClinVar one star: 14,819 ClinVar variants (7346 benign and 7473 pathogenic variants) that are assigned one star or above (meaning at least 1 laboratory (primarily clinical laboratories) have provided their rationale for variant assertion).

ClinVar 2 star: 2966 (1914 benign and 1052 pathogenic) ClinVar variants with two star status or above. These variants have concordant assertions from at least two independent laboratories.

ClinVar Oct 2015 to December 2016: This dataset contains 6949 (4093 benign and 2856 pathogenic) variants in ClinVar that were obtained from the variant_summary.txt file released in December 2016 after removing the variants that were present in the October 2015 data release.

ClinVar Sept 2016 to March 2017: This is a set of 3792 benign and 1310 pathogenic missense variants with one star or above ClinVar review status. These were obtained by filtering out the variants in the variant_summary.txt file in ClinVar from September 2016 from the variant_summary.txt files from March 2017 in ClinVar.

Oncogene variants: This dataset consists of 87 benign and 321 pathogenic variants in oncogenes as defined by genes having a high oncogene score and a low TSG score as described in [29].

Tumor suppressor gene variants: This dataset consists of 502 benign and 532 pathogenic variants in tumor suppressor genes as defined by genes having a high TSG score and a low oncogene score as described in [29].

Dominant: This dataset contains variants in genes that were associated with dominant mode of inheritance as determined by both[37] and[38]. There were 480 benign and 1591 pathogenic variants in this dataset.

Recessive: This dataset contains variants in genes that were associated with recessive mode of inheritance as determined by both [37] and [38]. There were 1169 benign and 1429 pathogenic variants in this dataset.

REVEL testset: This is the test dataset that contained ClinVar variants (Test Data 2) as described in [19].

MetaSVM/LR testset: This dataset consisted of 12496 (6275 benign and 6221 pathogenic) ClinVar variants (with one or more review status in ClinVar) which did not include the variants used in the training sets of MetaSVM/LR. predictSNPdsel: This is a benchmark dataset as described in[31]. It does not contain CADD training data.

Varibenchselected: This is a highly unbalanced dataset as described in [31]. According to the authors, more than 98% of all proteins in this dataset contain variants that are either “pathogenic” or “neutral”.

Balanced dataset: This dataset contained 4192 variants in ClinVar (one star or above status) with each gene having the same number of benign and pathogenic variants.

## Additional material

**Additional Figure 1:** Variability in performance of algorithms shown in each panel across all analyzed datasets. Performance is measured using AUC and depicted along the y axis. The algorithms are sorted by their performance and the datasets the color coded as outlined in the legend. The horizontal dotted red line shows an AUC of 0.8.

**Additional Figure 2:** Performance analysis of algorithms. The AUC of a ROC are plotted for 25 algorithms. Vertical dotted line indicates a AUC of 0.9 and 99% confidence intervals for each AUC are shown. AUCs of the algorithms across four different datasets (represented in panels) to address type II circularity as described in text.

**Additional Table 1:** Description of algorithms used in the analyses.

**Additional Table 2:** Number of variants and their review statuses for which majority of algorithm assertion was opposite to that in ClinVar.

**Additional Table 3:** Concordance among different combination of algorithms.

**Additional datasets:**

Additional data 1: ClinVar dataset of 14819 variants from September 2016

Additional data 2: True and false concordances for combinations of 2 algorithms

Additional data 3: True and false concordances for combinations of 3 algorithms

Additional data 4: True and false concordances for combinations of 4 algorithms

Additional data 5: True and false concordances for combinations of 5 algorithms

Additional data 6: Dataset with variants in Oncogene and TSG

Additional data 7: Dataset with variants in genes associated with dominant or recessive inheritance.

Additional data 8: Variants in ClinVar txt files (October 2015 to December 2016)

Additional data 9: Variant in ClinVar txt files (September 2016 to March 2017)

Additional data 10: No training variants MetaSVM/LR

Additional data 11: Dataset with variants in genes with different missense Z cutoff

Additional data 12: Balanced dataset

Additional data 13: Benchmark dataset: predictSNPsel

Additional data 14: Benchmark dataset: varibenchsel

Additional data 15: Compiled AUCs of algorithms for different datasets.

Additional data 16: Data after hierarchical clustering using pvclust.

Additional data 17: Code used for generating Figures and some of the additional data. (Also available at <u>http://rpubs.com/thisisrg/supp_code_17</u>)

## Acknowledgements

We grateful to the following people who provided us with algorithm scores for the variants and data used in the manuscript: Vikas Pejavar and Pedrag Radiovojac for providing us with some of the Mutpred scores, Panos Katsonis and Olivier Lichtarge for providing us with hEAt and EA scores. Weiva Sieh and Joseph Rothstein for providing the REVEL test data set. This work was supported by NHGRI 5 U01 HG007436-03 grant to SEP. Competing interests: SEP serves on the scientific advisory board of Baylor Genetics Laboratory.

